# The evolutionary age-range size relationship is modulated by insularity and dispersal in plants and animals

**DOI:** 10.1101/2023.11.11.566377

**Authors:** Adriana Alzate, Roberto Rozzi, Julian A. Velasco, D. Ross Robertson, Alexander Zizka, Joseph A. Tobias, Adrian Hill, Christine D. Bacon, Thijs Janzen, Loïc Pellissier, Fons van der Plas, James Rosindell, Renske E. Onstein

## Abstract

Earth is home to millions of plant and animal species, with more than 40 thousand species facing extinction worldwide (Diaz et al. 2019). Species’ range size is particularly important in this context because it influences extinction risk (Purvis et al. 2000, Gaston & Fuller 2009), but the causes underlying the wide natural variation in range size remain poorly known. Here, we investigate how evolutionary age is related to present-day range size for over 25,000 species of mammals, birds, reptiles, amphibians, reef fishes, and plants. We show that, on average, older species have significantly larger ranges, but the effect of age on range size is modulated by clade, geographical context and dispersal ability. Specifically, age does not affect range size for island species, because islands limit dispersal and hence range size, regardless of species age. Furthermore, species from clades with high dispersal capabilities obtain large ranges faster, thereby further neutralizing the relationship between age and range size. Our results can help supporting global conservation priorities, by showing that species that are young, occupy islands, and/or are dispersal limited often have small ranges and therefore increased extinction risk.

## Main

What drives species’ range size? This is a long-standing question in macroecology and biogeography with strong implications for conservation. Species with narrow geographical distributions face a higher risk of extinction compared to widespread species (Jablonski 1986, Purvis et al. 2000). This is because narrow-ranged species have lower overall abundance, and an increased likelihood that all the localities where they occur experience synchronous disturbances, resulting in local extinctions (Gaston & Fuller 2009). Consequently, understanding how ecological and evolutionary factors affect species’ geographical distributions can provide valuable insights for setting global conservation priorities.

One of the potential drivers of species range size variation is species ‘evolutionary age’ (age hereafter). Older species are expected to have larger geographical ranges than younger species because they have had more time for range expansion after speciation (the ‘age and area model’; Willis 1922). Despite the expected link between age and range size, previous studies have provided conflicting results (Webb & Gaston 2000, Böhning-Gaese et al. 2006, Mora et al. 2012, Swaegers et al. 2014, Hodge & Bellwood 2015). While support for a positive effect of age on range size was generally observed in fossil data for molluscs and trilobites (Jablonski 1987, Miller 1997), data for extant organisms show a more complex picture (Table S1). These previous studies varied strongly in their extent (e.g., focusing on regional versus global ranges), taxonomic scope (e.g., genus, family, order), and methodological aspects (e.g., method to quantify range size). Furthermore, various factors other than species age may influence range size, and this limits our understanding of a possible general effect of age on range size (Webb & Gaston 2000).

Species range size is the result of a variety of processes, including the rate by which species expand their range, the variety of habitat types that a species can persist in (‘niche breadth’), and colonization-extinction dynamics (Gaston 2003). Such processes can be influenced by a range of ecological, evolutionary and geographical factors that may subsequently obscure the age-range size relationship. For example, factors affecting the rate of range expansion (e.g., dispersal ability) and the maximum potential range that a species can attain (e.g., geographical context) may limit the effect of age on range size (Rayfield et al. 2023): species living on small, isolated islands may attain narrow ranges because of geographical constraints, even when they are old. Similarly, more dispersive species may attain larger ranges than less dispersive species (Alzate & Onstein 2022) because of faster range expansion, so young dispersive species may have larger range sizes than expected based on age only. Moreover, such range-size dynamics are likely species and lineage-specific, with less dispersive or wide-spread lineages (e.g., amphibians) showing more pronounced age-range size relationships than more dispersive or wide-ranging lineages (e.g., marine mammals) (Webb & Gaston 2000). Disentangling the effect of age on range size therefore requires accounting for the ecological and geographical factors that may have modulated range size through time.

Here, we investigated the relationship between species’ age and range size for over 25,000 species from the Tree of Life, including marine and terrestrial mammals, birds, amphibians, reptiles, reef fishes and palms. We hypothesize that i) evolutionary age has an overall positive effect on range size, but the strength of the effect depends on the clade, geographical context and dispersal abilities of the species; ii) the relationship between evolutionary age and range size is more pronounced when compared between broad taxonomic clades (e.g., orders) than within narrow taxonomic clades (e.g., families), because variability in range size and/or age decreases for narrow clades composed of closely related species that share more similar ages, distributions, and dispersal-related traits; iii) the relationship between evolutionary age and range size is absent on islands, because the maximum range size of endemic island species is geographically constrained, rather than age-constrained, and iv) the relationship between age and range size is absent for more dispersive species, because dispersal allows for faster range expansion than expected given species age, leading to dispersive species more quickly attaining maximum range sizes than less dispersive species. The results of our analyses show that age has an overall positive effect on species range sizes, but this relationship is clade-specific, constrained by the geographical context, and modulated by the dispersal abilities of the species.

## Results

### Evolutionary age positively affects range size

We first investigated whether age had a positive relationship with range size for 25,616 species from lineages of seven major groups of animals and plants, using linear mixed-effects models, and including taxonomic group as a random effect on slope and intercept. We found that evolutionary age had an overall significant positive effect on range size (Z = 0.10, SE = 0.02, df = 4.46, t = 3.97, p = 0.013; Fig. 1a), but the strength of the relationship varied among taxonomic groups (Fig. 1b-h, Table S2). For instance, reef fish, reptiles and amphibians show the strongest effect of age on range size, whereas terrestrial mammals show the least strong effect (Fig.1). These results were robust to outliers (Fig. S1, Table S3). Next, we investigated whether the strength of the age-range size relationship was clade-dependent, by running separate linear models at three taxonomic levels: broad taxonomic level (marine mammals, terrestrial mammals, amphibians, reptiles, reef fishes, palms and birds), order level (108 orders), and family level (418 families). To test the overall effect of age on range size, we ran three meta-analyses including all the individual relationships at the three taxonomic levels. We found taxon-dependent relationships between age and range size at the three taxonomic levels examined. At a broad taxonomic level, the effect of evolutionary age on range size was positive for the majority of the groups, with the exception of marine and terrestrial mammals (Fig. 2a, Table S3). At the order level, 70 out of 86 orders showed neutral relationships, nine were positive and seven were negative (Fig. 2b, Table S4). At the family level, 346 out of 410 families showed neutral relationships, five showed positive and 29 showed negative relationships (Fig. 2c, Table S4). The meta-analysis confirmed the overall positive effect of evolutionary age on range size at the broad taxonomic level (K = 7, Estimate = 0.08 [0.03 - 0.13], test for heterogeneity: Q = 48.26, df = 6, p < 0.0001), and at the family level (K = 410, Estimate = 0.03 [0.001 - 0.06], test for heterogeneity: Q = 1522.88, df = 409, p < 0.0001). However, there was a neutral effect at the order level (K = 86, Estimate = 0.05 [-0.003 - 0.100], test for heterogeneity: Q = 379.98, df = 85, p < 0.0001). These results indicate that evolutionary age-range size relationships are widespread and positive at broad taxonomic levels, but become weaker and less consistent at lower taxonomic levels, possibly due to lower variability in range size or age for groups composed of closely related species that share similar ages, distributions, and dispersal-related traits.

**Fig. 1.**
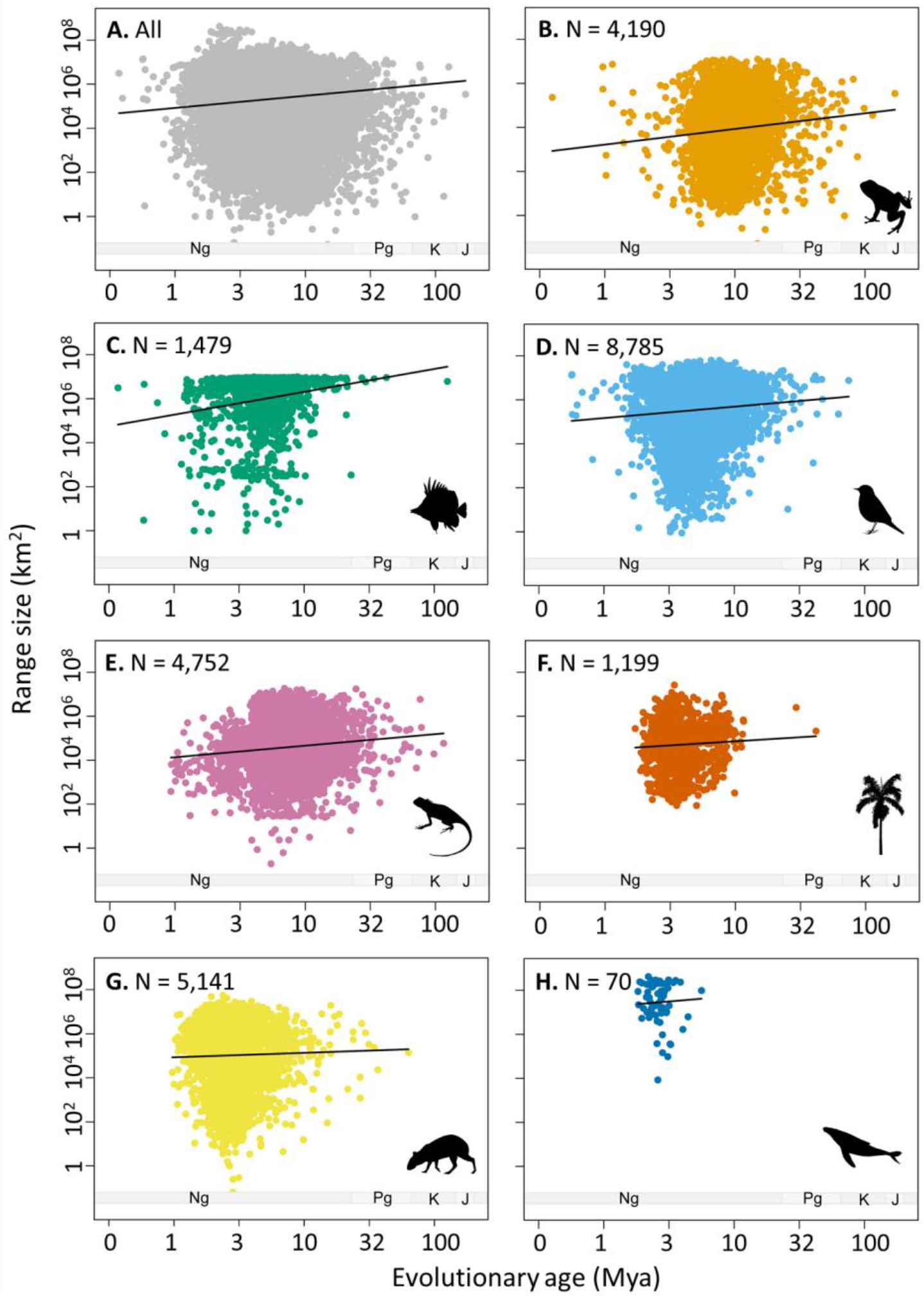
Relationships between evolutionary age (Mya = million years ago) and range size for a) all c. 25,000 species, and b-h) separated for the six taxonomic groups: b) amphibians, c) reef fishes, d) birds, e) squamates, f) palms, g) terrestrial mammals, and h) marine mammals. The black line in a) denotes the overall positive relationship between evolutionary age and range size and in b–h) it denotes the relationship for each taxonomic group (the taxon-specific slopes of the model). Geological periods are denoted as Ng = Neogene, Pg = Paleogene, K = Cretaceous, and J = Jurassic. Data were log-transformed for analyses and plotting.

**Fig. 2.**
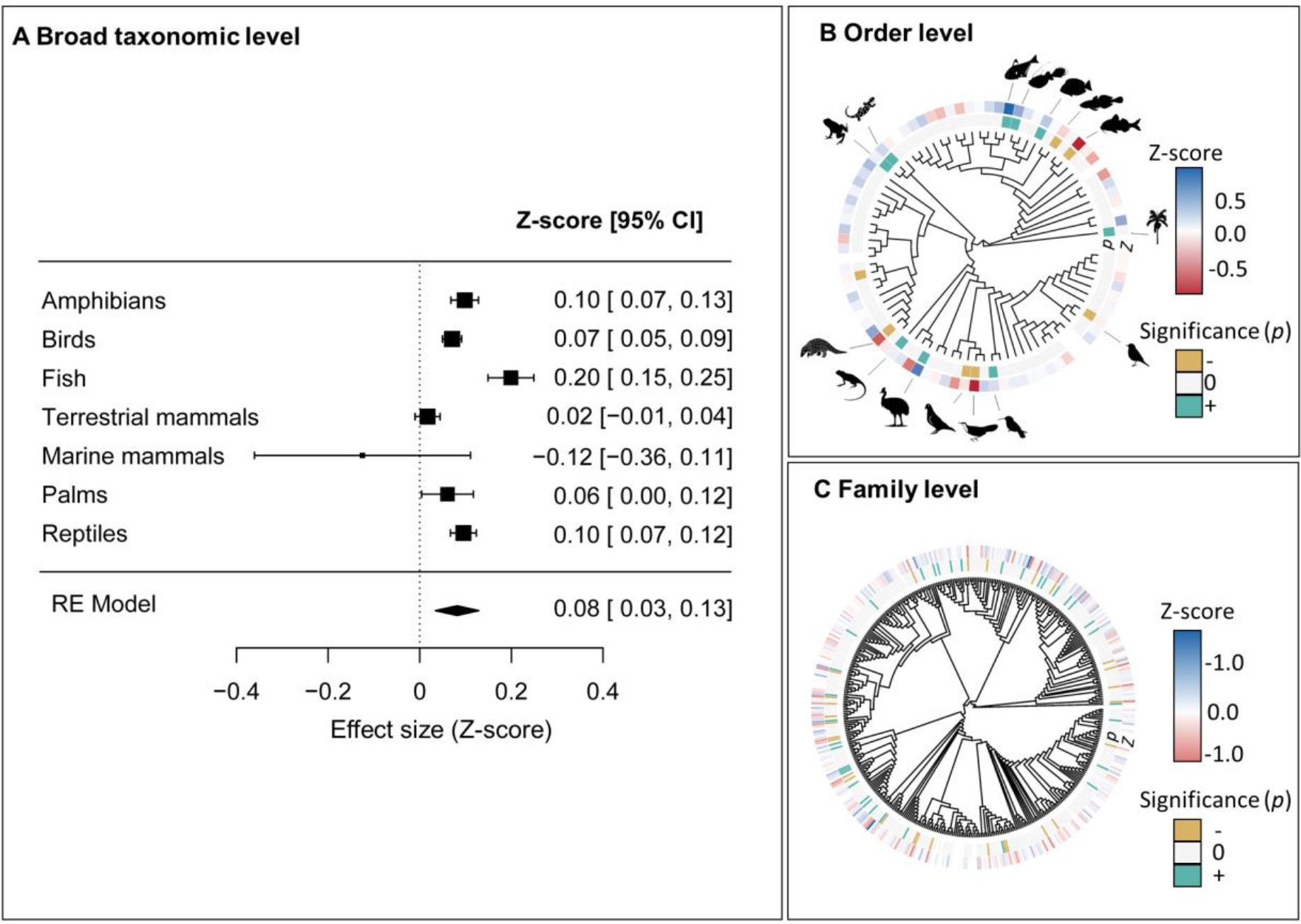
a) Forest plot of the effect of evolutionary age on range size for the seven broad taxonomic groups, including the overall effect from a random effects (RE) model. b) Effect of evolutionary age on range size at the order level. c) Effect of evolutionary age on range size at the family level. Outer arc colours denote the strength (Z-score) of the relationship between evolutionary age and range size, and inner arc colours denote the significance. Icons indicate orders with significant relationships. CI = Confidence Interval.

### Geographical constraints on the relationship between evolutionary age and range size

To test whether the effect of age on range depends on geographical context (species restricted vs. not restricted to islands), we ran two individual linear mixed-effect models: one for the subset of species restricted to islands (N = 3,573) and one for the species not restricted to islands (N = 22,113). We included the broad taxonomic group as a random effect. We found that the range size a species can attain is constrained by geography, with species restricted to islands attaining on average smaller ranges than species not restricted to islands (Fig. 3, Fig. S2). Moreover, the relationship between age and range size was modulated by that geographical context. Species that were not restricted to islands exhibited a positive effect of evolutionary age on range size (Estimate = 0.11 [0.06-0.16], SE = 0.02, df = 4.86, p = 0.007; Fig. 3a, Table S7), whereas species restricted to islands did not show an effect of evolutionary age on range size (Estimate = 0.03 [-0.02 - 0.08], SE = 0.02, df = 3099, p = 0.18; Fig. 3b, Table S8). This pattern was consistent across all taxonomic groups, except for terrestrial mammals, where there was no significant relationship between age and range size, irrespective of geographical context. These results suggest that the range size of island species is constrained by the maximum range a species can attain and likely by factors hindering range expansion, thereby uncoupling the relationship between species age and range size.

**Fig. 3.**
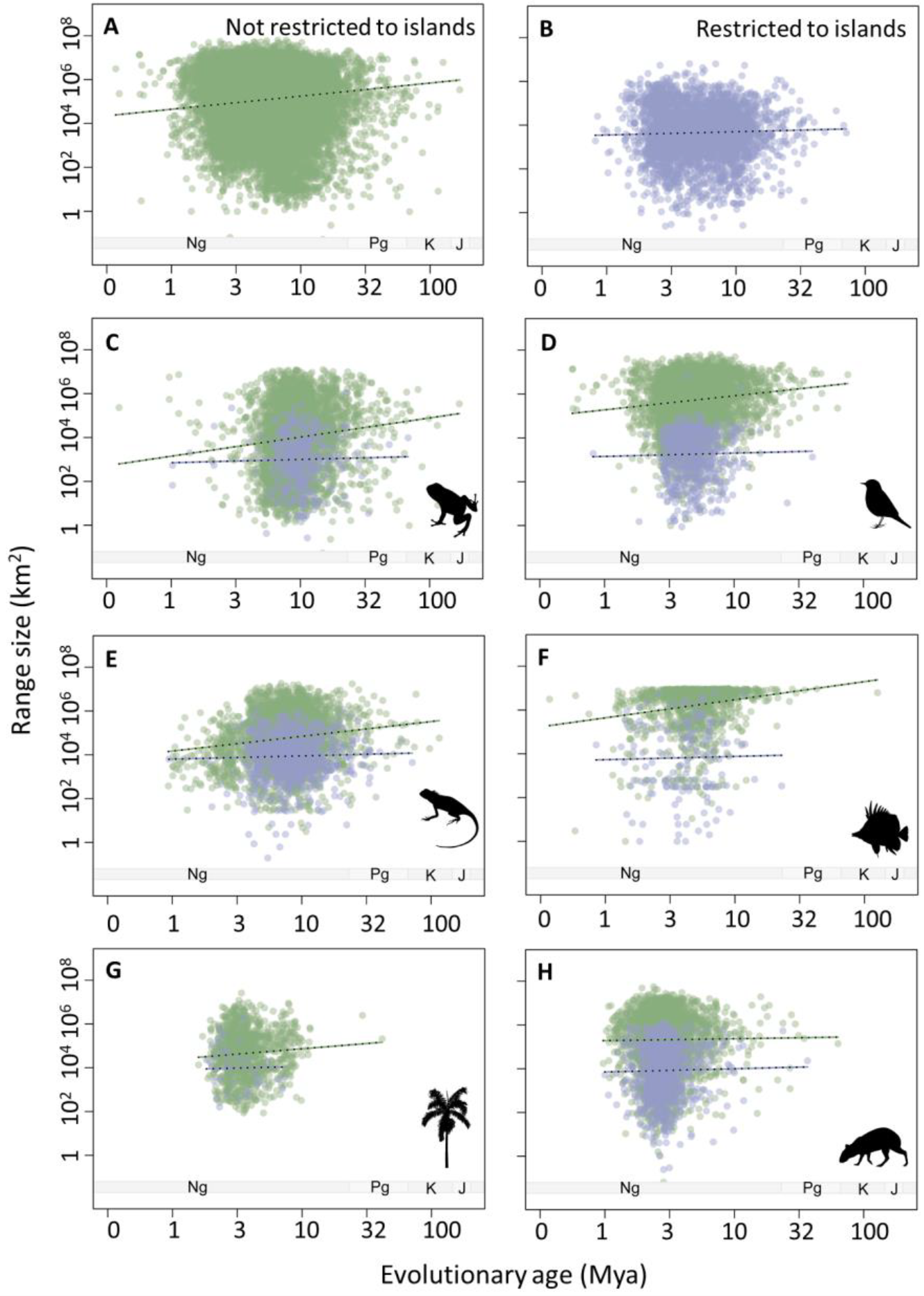
Relationship between evolutionary age (Mya = million years ago) and range size for a) all species that are not restricted to islands, b) all species restricted to islands, and c-h) separated for the six taxonomic groups: c) amphibians, d) birds, e) squamates, f) reef fishes, g) palms, and h) terrestrial mammals. The dashed lines in a-b) denote the overall positive effect of evolutionary age on range size and in c-h) the relationship for each taxonomic group (the random slopes of the model) for species not restricted to islands (green) and restricted to islands (purple). Geological periods are denoted as Ng = Neogene, Pg = Paleogene, K = Cretaceous, and J = Jurassic. Data were log-transformed for analyses and plotting.

### Dispersal abilities modulate the evolutionary age-range size relationship

We investigated whether the effect of age on range size varied with the dispersal abilities of species, by running individual linear mixed-effects models for each taxonomic group. We included dispersal-related traits and evolutionary age as interacting and fixed effects. We included family, continent/region (Americas, Asia, Africa, Europe, Australia, Greater Caribbean, Tropical Eastern Pacific) and geographical context (restricted or not restricted to islands) as random effects. For each group, we used different proxies to quantify dispersal, such as body size, hand-wing index and fruit size (see methods). We found that the strength of the relationship between age and range size depended on the dispersal abilities of the species, but not for all taxonomic groups (Fig. 4, Table S9). For example, reef fishes with higher dispersal abilities (larger body sizes) showed a neutral age-range size relationship, whereas reef fishes with lower dispersal abilities showed a positive relationship. In contrast, for marine mammals and palms, the effect of age on range size only became apparent for species with higher (larger body sizes or larger fruit sizes) instead of lower dispersal abilities. For amphibians, reptiles and birds, the relationship between age and range size was not affected by dispersal ability, as indicated by a non-significant interaction effect between age and dispersal. However, for reptiles and amphibians, there was a trend for more dispersive (i.e., larger-bodied) species to be less affected by the age effect on range size than for less dispersive (i.e., smaller-bodied) species. Together, the positive relationship between species age and range size is more prominent in taxa or species with lower dispersal abilities.

**Fig. 4.**
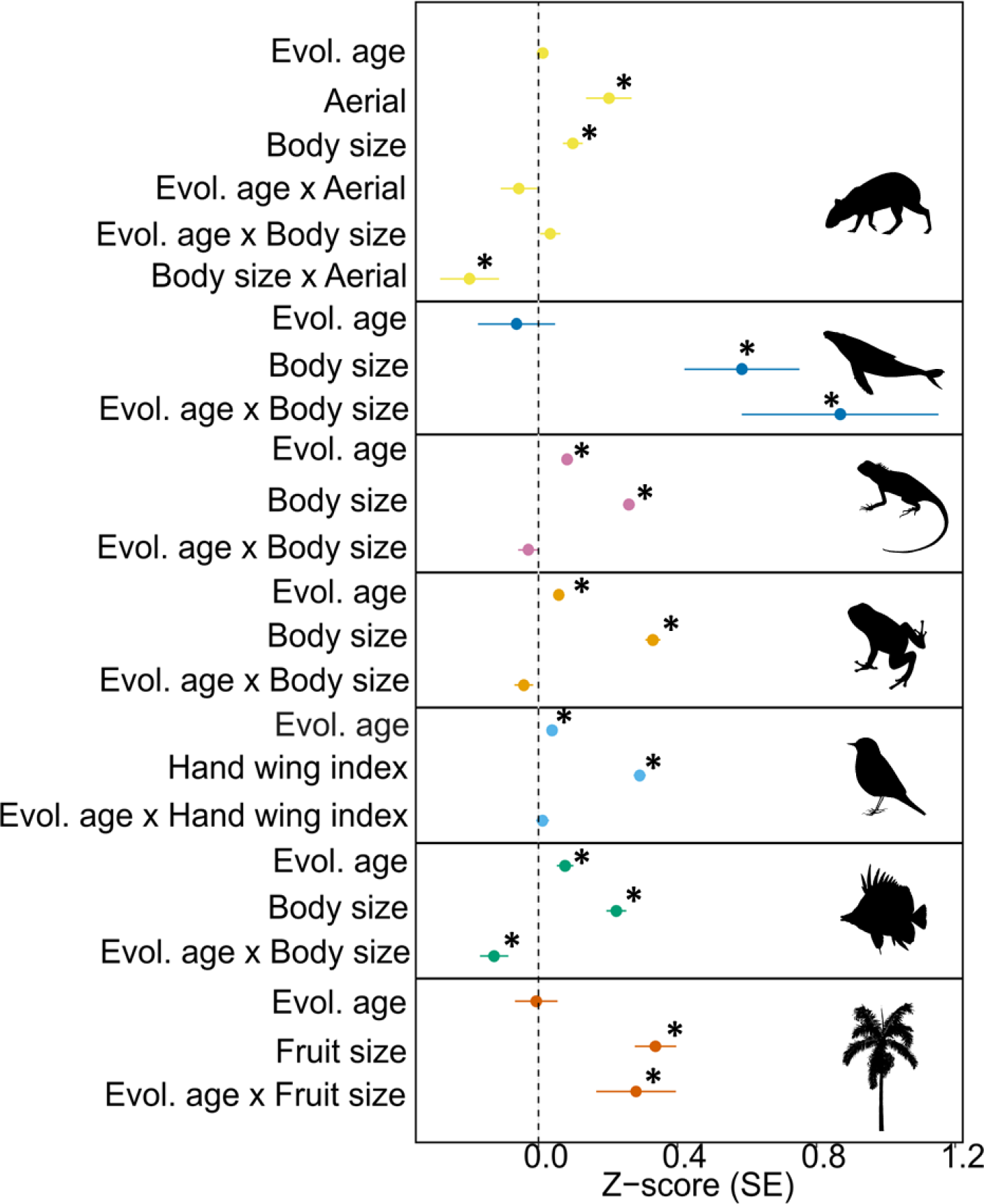
Dispersal ability modulates the relationship between evolutionary age and range size. Effect of evolutionary age and dispersal-related traits on species range size for terrestrial and marine mammals, squamates, amphibians, birds, reef fish and palms. We considered dispersal-related traits ‘aerial dispersal’, ‘body size’, hand-wing index’ and ‘fruit size’ as proxies of dispersal. Asterisks denote significant relationships. Evol. age = evolutionary age; ‘x’ refers to an interactive effect. Points indicate the effect size (Z-score) and the error bars the standard error (SE). Some data points lack error bars because of very small SE values.

Overall, our results showed that older species with higher dispersal abilities that are not restricted to islands attain larger range sizes than their younger counterparts that have lower dispersal abilities and/or that are restricted to islands. Thus, to understand and predict current and future species range sizes, we should consider the macroevolutionary time for range expansion, the geographical context, and the dispersal abilities of the species. Furthermore, we showed that the age-range relationships were highly variable when examined for single orders or families, which may provide an explanation for the mixed (positive, neutral, or negative) age-range size relationships detected in previous studies (Table S7-9). Many of the studies to date have focused on few species at narrow taxonomic levels (e.g., within a single genus or family) or studied the relationship in a confined geographical area, such as islands (Wollenberg et al. 2011, Pepke et al. 2019) or specific biomes, such as the South African fynbos (Schurr et al. 2007) and the Brazilian Atlantic forest (Leão et al. 2020). Consequently, the variability in the age-range size relationship we observed in this study, particularly within single orders or families (Fig. 2), is consistent with the variety of age-range size relationships detected in previous studies. We show that this variability can, at least partly, be explained by variation in dispersal ability, geographical context and scope, and taxonomic group and level.

The evolutionary age-range relationship was mostly non-significant for species restricted to islands. Geographical constraints exerted by islands influence the maximum range species can attain, and thereby modulate the relationship between evolutionary age and range size. Several factors may explain why species on islands have, on average, smaller ranges than expected by their evolutionary age. Range expansion from islands is limited due to physical barriers (oceans) to dispersal for most taxa. Theoretical models have also shown that habitat fragmentation has a stronger effect on the range expansion of species with limited dispersal abilities than for more dispersive ones (Alzate et al. 2019a). Furthermore, the commonly observed evolutionary loss of species’ dispersal abilities on islands can impact range expansion and speciation (Waters et al. 2020, Fernández-Palacios et al. 2021). However, species restricted to islands have similar average evolutionary ages to species not restricted to islands (Fig. S3). Species restricted to islands may therefore often be relics of species or lineages that were once more widespread but have experienced extinctions elsewhere, as shown in birds (Oswald et al. 2023). This would disconnect current range size from species age. Furthermore, species restricted to islands may be more likely to go extinct before expanding their ranges (Simberloff 2000). Our results suggest that islands act as eco-evolutionary ‘traps’, limiting range expansions due to dispersal barriers, local extinctions, and the subsequent evolutionary loss of dispersal abilities, thereby increasing the risk of extinction due to small range sizes.

Dispersal ability can modulate the evolutionary age-range size relationship in two ways. On the one hand, high dispersal ability allows species to expand and reach large range sizes rapidly, possibly reducing the influence of age on range expansion and size. On the other hand, high dispersal ability prevents speciation and, thereby, range splitting, by maintaining gene flow among populations (Mayr 1963, Yamaguchi 2022). Indeed, dispersal ability positively affects range size (Alzate & Onstein 2022), and we found that for all clades examined and for species of similar age, those with higher dispersal abilities had larger ranges than species with lower dispersal abilities (Fig.4). This suggests that dispersal modulates how fast species expand their ranges after speciation, and may obscure the relationship between age and range size. Range expansion is expected to happen more gradually for less dispersive species or clades, leading to an increased effect of age on range expansion and size, as observed here in reef fishes, amphibians and reptiles. Interestingly, despite their comparatively old evolutionary ages, amphibians exhibited narrow geographical ranges, reflecting their limited dispersal capacities (Jreidini & Green 2023), specialized habitat requirements, and slow niche evolution (Rolland et al. 2018).

Interestingly, the positive effect of age on range size for highly dispersive palms and marine mammals contradicts our hypothesis that dispersive species attain maximum range sizes faster than less dispersive species, reducing the influence of age on range size. Possibly, historical and biological features of these groups explain this finding. For instance, palms with large fruits often rely on large-bodied fruit-eating and seed-dispersing animals (frugivores) for seed dispersal and range expansion (Onstein et al. 2017). Such large-bodied frugivores can move seeds over long distances in their often large home ranges (Pires et al. 2017). However, many large-bodied animals (megafauna) are now extinct, limiting dispersal of (old) palm species with large fruits, thus potentially leading to local extinctions and range reductions (Onstein et al. 2018, Méndez et al. 2022), and the recovery of the age-range size relationship. For marine mammals, the energetic constraints derived from large body sizes, including both a need to forage in larger areas and larger fasting capacities, allow more dispersive marine mammals to attain larger global distributions (Boyd 2004). Moreover, migratory behaviour, the occurrence of which positively correlates with body size in marine mammals (Boyd 2004), can occasionally lead to the diversification of lineages in new environments (Winker 2000), likely accentuating the effect of age on range size.

While we observed an overall positive relationship between species’ evolutionary age and their geographical range size, there are several factors that may have limited the strength of this relationship. First, species’ geographical ranges are dynamic over time (Pigot et al. 2012). Historical events such as paleoenvironmenal changes, tectonics, vicariance and habitat shifts (Wallace 1876, Simpson 1964, Brooks and McLennan 1991, Hagen et al. 2021), have played important roles in shaping current biodiversity and distribution patterns through differential speciation and (local) extinction regimes. For example, the impact of the last glacial period, notably in regions such as Eurasia, caused habitat contractions and pushed species to lower latitudes or confined them to smaller local glacial refugia (Price et al. 1997). During vicariance events, newly formed species may display various range sizes and range shapes, because ancestral species’ ranges are split asymmetrically (Anacker & Strauss 2014). Second, differential speciation-extinction patterns (Schwartz & Simberloff 2001) may lead to different species’ evolutionary age-range size relationships. A high speciation rate results in many young lineages and species with relatively small ranges (Alzate et al. 2019a, Leão et al. 2020). Furthermore, taxonomic classification biases, stemming from lumping or splitting methods, can impact species-level phylogenies and hence evolutionary age estimates (Schwartz & Simberloff 2001, Agapow et al. 2004, Faurby et al. 2015). The detection of recently diverged or near-extinct species, as well as the accurate estimation of range size, is impacted not only by extinction events but also by geographic sampling biases (Pigot et al. 2012, Gaston & Fuller 2009; Blackburn & Gaston 1998). Finally, human activities, as evidenced by significant extinctions during the Quaternary, particularly affecting large-bodied mammals (Faurby & Svenning 2015, Rozzi et al 2023), may have obscured the relationship between evolutionary age and range size, due to range contraction and extinctions (Borges et al. 2019, Pacifici et al. 2020). All these factors underline the intricate relationship between historical events and the dynamic nature of species’ ranges over time, and may explain why even though there are generally positive relationships between species age and range size, these relationships are not always strong or pronounced.

Our findings emphasize the complex interplay between evolutionary age, intrinsic dispersal abilities, and environmental/historical contexts, highlighting the need to consider all those factors in combination when elucidating causes of contemporary range sizes and biodiversity patterns. Our results offer new perspectives for shaping global conservation priorities, emphasizing the elevated risk of extinction for species restricted to islands and those of recent evolutionary origin with limited dispersal abilities, given their restricted range sizes.

## Methods

### Data collection

We collated range size data for 26,053 species (Fig. S4). We obtained distribution maps from the International Union for Conservation of Nature’s Red List of Threatened Species (IUCN) for amphibians, marine mammals and terrestrial mammals, and from Roll et al. (2017) for reptiles. For birds, we obtained range size data from the AVONET database (Tobias et al. 2022) and for palms from Hill et al. (2023). For reef-associated bony fishes (class ‘Actinopterygii’), we obtained occurrence data from Robertson & van Tassell (2019) and Robertson & Allen (2015). We documented and/or approximated range size as the extent of occurrence (EOO) within a projection that accounts for the curvature of the Earth.

We collated data on the evolutionary age for 25,616 species (Fig. S4). We obtained 100 phylogenetic trees for birds (Jetz et al. 2012), squamates (Tonini et al., 2016), amphibians (Jetz & Pyron, 2018) and mammals (Upham et al. 2019) from ‘Vertlife.org’. We obtained 100 phylogenetic trees for fish from the ‘Fish Tree of Life’ (Rabosky et al., 2018), and for palms from Faurby et al. (2016). We estimated the mean evolutionary age per species as the average branch length of the 100 phylogenetic trees terminal nodes (i.e., species or tips).

We collated data on species dispersal abilities for 25,449 species. For mammals, we obtained data on body size and flight ability from Phylacine 1.2.1 (Faurby et al. 2018, Faurby et al. 2019). For birds, we obtained data on hand-wing Index (HWI) from AVONET (Tobias et al. 2022). For amphibians, we obtained body size data from AmphiBIO (Oliveira et al., 2017) and for reptiles from Feldman et al. (2016). For reef fishes, we obtained body size data from Alzate et. al (2019b), Robertson & van Tassell (2019) and Robertson & Allen (2015). For palms, we obtained fruit size data from Kissling et al. (2019).

We collated data on insularity (landmasses smaller than Australia) for 25,692 species. For terrestrial mammals, we obtained data on island endemicity from Phylacine 1.2.1 (Faurby et al. 2018, Faurby et al. 2019). We consider bird species as “restricted to islands” when they were reported to be 100% associated with islands in Sheard et al. (2020). For amphibians, we obtained information on insularity by overlaying species distribution maps with a shapefile of islands of the world. For reptiles, we obtained information on geographical context (“restricted to islands” vs. “not restricted to islands”) from Meiri (2019), and for reef fishes from Alzate and colleagues (2019), Robertson & van Tassell (2019) and Robertson & Allen (2015). We classified palm species as restricted or not restricted to islands based on Cassia-Silva et al. (2021).

### Data analyses

To examine the overall effect of evolutionary age on range size we ran a linear mixed model including 25,616 species from 7 clades. We used the function ‘lmer’ from the ‘lme4’ R package (Bates et al. 2015). We included random slopes for each taxonomic group.

To examine how the evolutionary age-range size relationship varied among clades, we ran linear models at three taxonomic levels: at the broad taxonomic level (reef fish, birds, terrestrial mammals, marine mammals, amphibians, squamates and palms), at the order level (88 orders), at the family level (418 families). To test the overall effect of evolutionary age on range size based on all individual relationships, we performed a meta-analysis for each taxonomic level.

We fitted a random effects model including the Z-score values with their corresponding standard errors, using the function ‘rma’ from the R package ‘metafor’ (Viechtbauer 2010).

We tested the effect of geographical constraints on the relationship between evolutionary age and range size, as range expansion may be hindered for species restricted to island systems. Thus, we compared the effect of evolutionary age on range size for species that exclusively live on islands, to species that occur on the mainland (continents, or on continents and islands) by using a linear mixed-effects model with taxonomic group as a random slope. We used the function ‘lmer’ from the ‘lme4’ R package (Bates et al. 2015). Marine mammals were excluded from the analysis as too few species were restricted to islands.

To test whether the relationship between evolutionary age and range size depended on dispersal, we ran linear mixed-effects models for the six broad taxonomic levels (reef fish, birds, terrestrial mammals, marine mammals, amphibians, squamates and palms). We included evolutionary age and dispersal-related traits as fixed direct and interactive effects. Continents (Americas, Africa, Asia, Australia, Europe) or marine regions for reef fish (Greater Caribbean, Tropical Eastern Pacific), geographical context (restricted or not restricted to islands) and family, were included as random slopes. All dispersal-related traits were log-transformed and rescaled for the analyses. We accounted for geographical constraints and palaeogeographic history by including whether species are restricted to islands and the continent/region as random intercepts, except for marine mammals, which are all restricted to the open ocean and have global, circumtropical or circumtemperate distributions.

Evolutionary age and range size were log-transformed (log10) to meet linearity assumptions. All models were standardized using the function ‘standardize’ from the ‘arm’ R package (Gelman & Hill 2007). See supplementary materials for a more extensive description of the methods.

## Acknowledgements

This study was funded by iDiv through a German Research Foundation grant (DFG FZT 118, 202548816). AA received support from an individual postdoctoral grant from sDiv and a Marie Skłodowska-Curie postdoctoral grant (Grant No. 101061723), funded by the European Union’s Horizon 2020 research and innovation programme. CDB was supported by the Swedish Research Council (#2017-04980; #2022-03927) and the University of Gothenburg Strategic Research Area in Biodiversity and Ecosystem Services in a Changing Climate (BECC). JAV received support from a DGAPA/PAPIIT UNAM grant (IA201320. We also thank all data collectors and providers.

## Author contributions

Conceptualization: AA, JR, LP, REO; Methodology: AA, FvdP, REO; Formal analysis: AA; Data Curation: JAV, AH, CDB, JT, DRR, AA; Writing - Original Draft: AA; Writing - Review & Editing: AA, RR, FvdP, JAV, DRR, AH, CDB, JT, DRR, TJ, LP, JR, REO.

## Materials & Correspondence

Adriana Alzate

